# Vast population genetic diversity underlies the treatment dynamics of *ETV6-RUNX1* acute lymphoblastic leukemia

**DOI:** 10.1101/117614

**Authors:** Veronica Gonzalez-Pena, Matthew MacKay, Iwijn De Vlaminck, John Easton, Charles Gawad

## Abstract

Ensemble-averaged genome profiling of diagnostic samples suggests that acute leukemias harbor few somatic genetic alterations. We used single-cell exome and error-corrected sequencing to survey the genetic diversity underlying *ETV6-RUNX1* acute lymphoblastic leukemia (ALL) at high resolution. The survey uncovered a vast range of low-frequency genetic variants that were undetected in conventional bulk assays, including additional clone-specific “driver” RAS mutations. Single-cell exome sequencing revealed APOBEC mutagenesis to be important in disease initiation but not in progression and identified many more mutations per cell than previously found. Using this data, we created a branching model of *ETV6-RUNX1* ALL development that recapitulates the genetic features of patients. Exposure of leukemic populations to chemotherapy selected for specific clones in a dose-dependent manner. Together, these data have important implications for understanding the development and treatment response of childhood leukemia, and they provide a framework for using population genetics to deeply interrogate cancer clonal evolution.

**One-Sentence Summary:** APOBEC and replication-associated mutagenesis contribute to the development of ETV6-RUNX1 ALL, creating massive leukemic population genetic diversity that results in clonal differences in susceptibilities to chemotherapy.

## Introduction

Analogous to organismal evolution in an ecosystem, the environmental selection pressures, population size, and rate of genome modification are core determinants of tumor evolutionary dynamics^1^. Contemporary bulk sequencing methods that interrogate the ensemble-averaged mutational profiles of the genomes of thousands of cells predominantly identify those mutations present in the most dominant tumor subclones at diagnosis^2^. However, bulk sequencing strategies are limited in their ability to capture the full genetic diversity of a population, as they do not identify lower-frequency clones and variants. A deeper understanding of the genetic diversity within tumors is key to understanding tumor evolution, including that of malignant cell populations that may increase in size as the relative fitness of clones changes in response to new selection pressures as patients undergo treatment.

Our knowledge of the evolutionary dynamics of acute leukemias is largely derived from bulk sequencing studies, which have concluded that acute leukemias harbor minimal genetic complexity when compared to other malignant neoplasms^3,4^. *In utero* acquisition of an *ETV6-RUNX1* translocation has been identified as the most frequent initiating event of acute lymphoblastic leukemia (ALL)^5,6^, the most common childhood leukemia. However, an *ETV6-RUNX1* translocation is insufficient for leukemogenesis^7^; recent genomic studies have identified deletions of genes required for normal B-cell differentiation^8^ that harbor signatures of aberrant RAG recombinase activity as frequent cooperating lesions^9^. In addition, *ETV6-RUNX1* ALL cells always harbor somatic single-nucleotide variants (SNVs)^9,10^. By using single-cell genomics, we determined that most of the deletions occur before the acquisition of SNVs, which result in the outgrowth of co-dominant clonal populations^9^. In addition, analyses of the SNV sequence motifs revealed an important mutagenic role for aberrant cytosine deaminase activity by an APOBEC (apolipoprotein B mRNA-editing enzyme, catalytic polypeptide-like) protein^9,10^. As APOBEC proteins mediate innate defense against viral infections^11^, this finding supports the hypothesis that environmental exposure to viruses triggers the transforming mutagenesis^12^. ALL also undergoes clonal evolution between its initial diagnosis and relapse^13,14^. However, the magnitude of that evolution and the changes in clonal composition between diagnosis and relapse remains unknown.

Several fundamental questions regarding the genesis and treatment response of *ETV6-RUNX1* ALL could be more precisely addressed by studying ALL genetics on a population scale. For example, why are there co-dominant clonal populations, especially when some clones harbor known “driver” mutations? What is the total mutation burden across the population of malignant cells, and do the underlying mutational processes change over time? How does this population genetic diversity influence treatment response?

To address these questions, we first used single-cell exome sequencing and error-corrected sequencing to further dissect the intraclonal genetic diversity and the shift in mutational processes that occurred during *ETV6-RUNX1* ALL development. These analyses revealed evidence of massive population genetic diversity. Then, after patients began therapy, we examined the evolution of those leukemic clones by exposing samples to standard chemotherapy drugs. Together, our findings enhance our understanding of the development and treatment response of *ETV6-RUNX1* ALL at the single-cell level and have significant clinical implications.

## Results

### Clone-specific “driver” mutations identified by single-cell exome sequencing

To further characterize the genomic diversity of clones previously defined by segregating variants identified in the bulk sample to single cells^10^, we performed single-cell exome sequencing on 3 cells from each clone and on 3 normal cells from the same patient (Fig. 1A). We achieved a mean saturating coverage breadth of 82% of the target exome with 60 million reads, as compared to 95% coverage of the target exome in bulk samples (**Supplementary Fig. 1**). Using only the single cells, we called mutations by requiring at least 2 cells to have the same base change at the same genomic position. The initial clonal structure had 5 high-frequency clones, with one of the 2 largest clones harboring an E63K *KRAS* mutation, whereas we could not clearly identify transforming alterations in the other clones (Fig. 1A). We then identified a further 10 to 29 mutations per clone (Fig. 1B). Using the normal cells as a control, we identified a low false-variant call rate at 7 sites, possibly resulting from clonal mutations acquired in nonmalignant cells, amplification artifacts, or sequencing errors. We then compared the mutagenic base-change pattern of the earlier, higher-frequency mutations that were detected in bulk and were shared between clones to the later, clone-specific changes that were detected only with single-cell exome sequencing. Interestingly, the early mutations were strongly enriched for C-to-T and C-to-G changes with an associated APOBEC motif (T preceding C), whereas the most common later mutations were A-to-G, A-to-C, and C-to-T, with enrichment for G following C. That signature is most consistent with replication errors (Fig. 1C,D)^15^. We also found a *KRAS* G12S mutation that was confined to a less abundant clone, as well as another clone-specific *NRAS* G12D variant; both of these mutations were C-to-T changes (Fig. 1A). Because the 3 activating *RAS* mutations (G12R, G12S, E63K) were acquired in distinct clones and not sequentially, we hypothesized that they occurred over a relatively short time. We then proposed that more oncogenic mutations would be acquired slightly later than the dominant clones, resulting in their being present at even lower frequencies.

**Figure 1.**
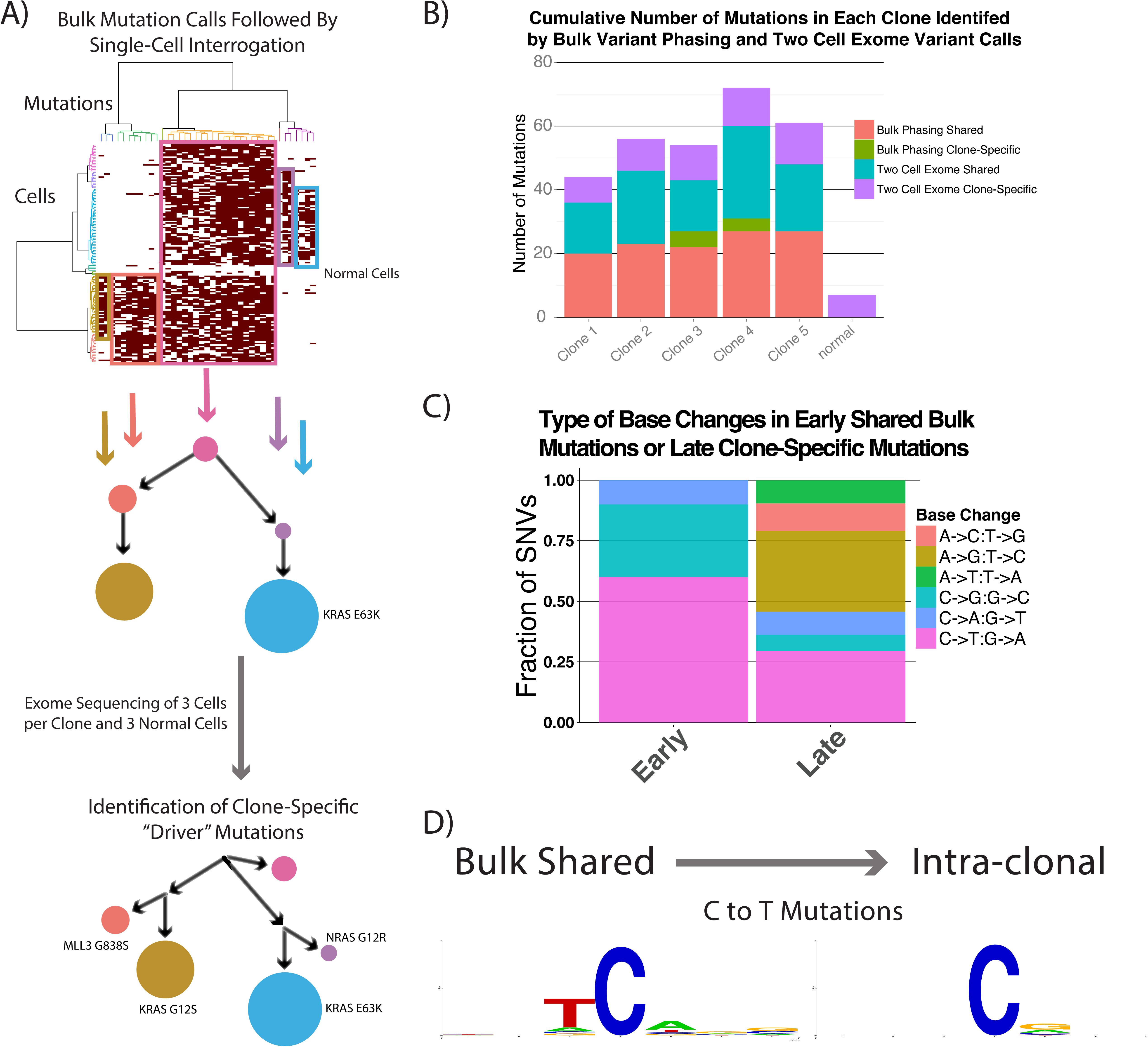
Identification of clone-specific “driver” mutations by using single-cell exome sequencing. (A) The clonal structure of an *ETV6-RUNX1* diagnostic patient sample that was identified by interrogating single cells for mutations first detected in the bulk sample was further resolved by calling mutations in the single cells alone. The clone-specific “driver” *RAS* mutations identified as possible causes of the clonal expansions are noted. (B) The number of new mutations identified in each clone using phasing of bulk mutations and 2-cell mutation calls. (C) Base substitution patterns seen in shared (early) and clone-specific (later) mutations. (D) The surrounding motifs in C-to-T mutations in early and late SNVs, showing that the strong APOBEC motif is only present in the early mutations.

### Deeper interrogation identifies massive population genetic diversity

To more deeply characterize oncogenic mutations in the bulk sample from the same patient, we performed error-corrected targeted sequencing of 50 mutational hotspots in ALL (**Supplementary Table 1**). We identified the same 3 *RAS* mutations (G12R, G12S, E63K) that were detected through bulk and single-cell exome sequencing, along with 2 additional known activating *KRAS* mutations (G12D, D119N) (Fig. 2A)^16–19^, all of which were C-to-T substitutions. We found no evidence of mutations in the other 37 genes that we examined using a Fisher's exact test cutoff of 0.01. The only benign lesion found in a *RAS* gene was a G12G mutation that was part of the dinucleotide G12D variant.

**Figure 2.**
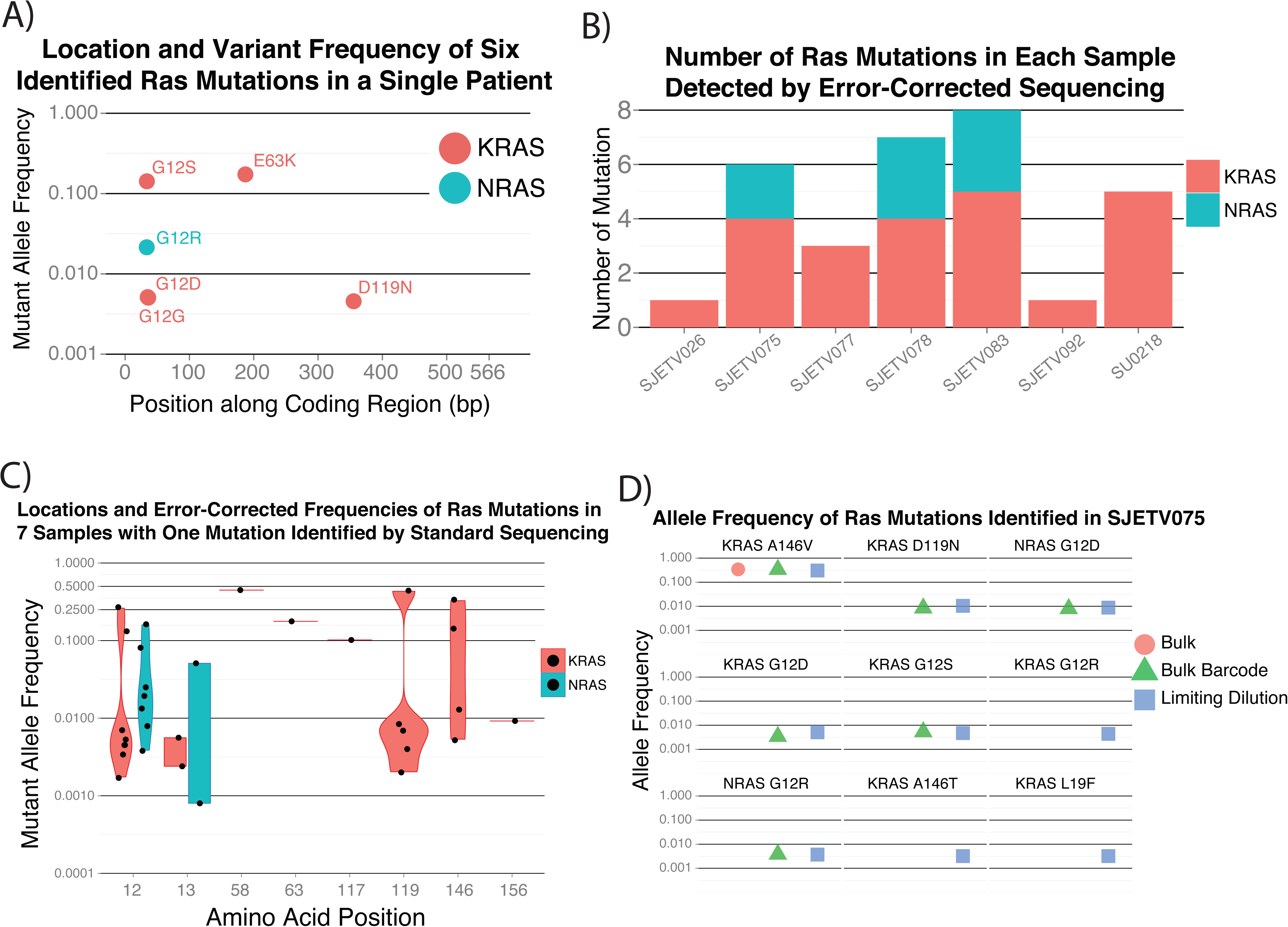
Evidence for large population genetic diversity in *ETV6-RUNX1* ALL. (A) Error-corrected sequencing confirmed 3 clone-specific activating *RAS* mutations and identified 2 additional lower-frequency activating mutations. (B) Subclonal *RAS* mutations were also common in a larger cohort in which each patient had one mutation identified in the bulk sample but had a median of 5 activating *RAS* mutations. (C) The allele frequency distributions of *RAS* mutations show no evidence of preferential selection of specific amino acid changes. (D) The increased sensitivity of mutation detection with limiting dilution identifies additional activating *RAS* mutations.

To evaluate whether these findings were applicable to a wider range of *ETV6-RUNX1* leukemias, we performed error-corrected sequencing on samples from 6 patients with a single *RAS* mutation identified by bulk sequencing and on 6 samples from patients with no known *RAS* mutations. We detected no *RAS* mutations in the latter 6 samples; however, we found a median of 5 activating *RAS* mutations in the 6 patients for whom a mutation was detected in the bulk samples (Fig. 2B). On examining the additional *RAS* variants more closely, we found the mutations to be clustered at known *RAS* mutational hotspots, namely *KRAS* codons 12, 13, 119, and 146 and NRAS codons 12 and 13. There was no clear propensity for variants at specific codons to be subclonal, supporting the assertion that the timing of the acquisition determined whether a mutation became dominant, as opposed to the change in RAS signaling activity due to the specific amino acid change (Fig. 3C).

**Figure 3.**
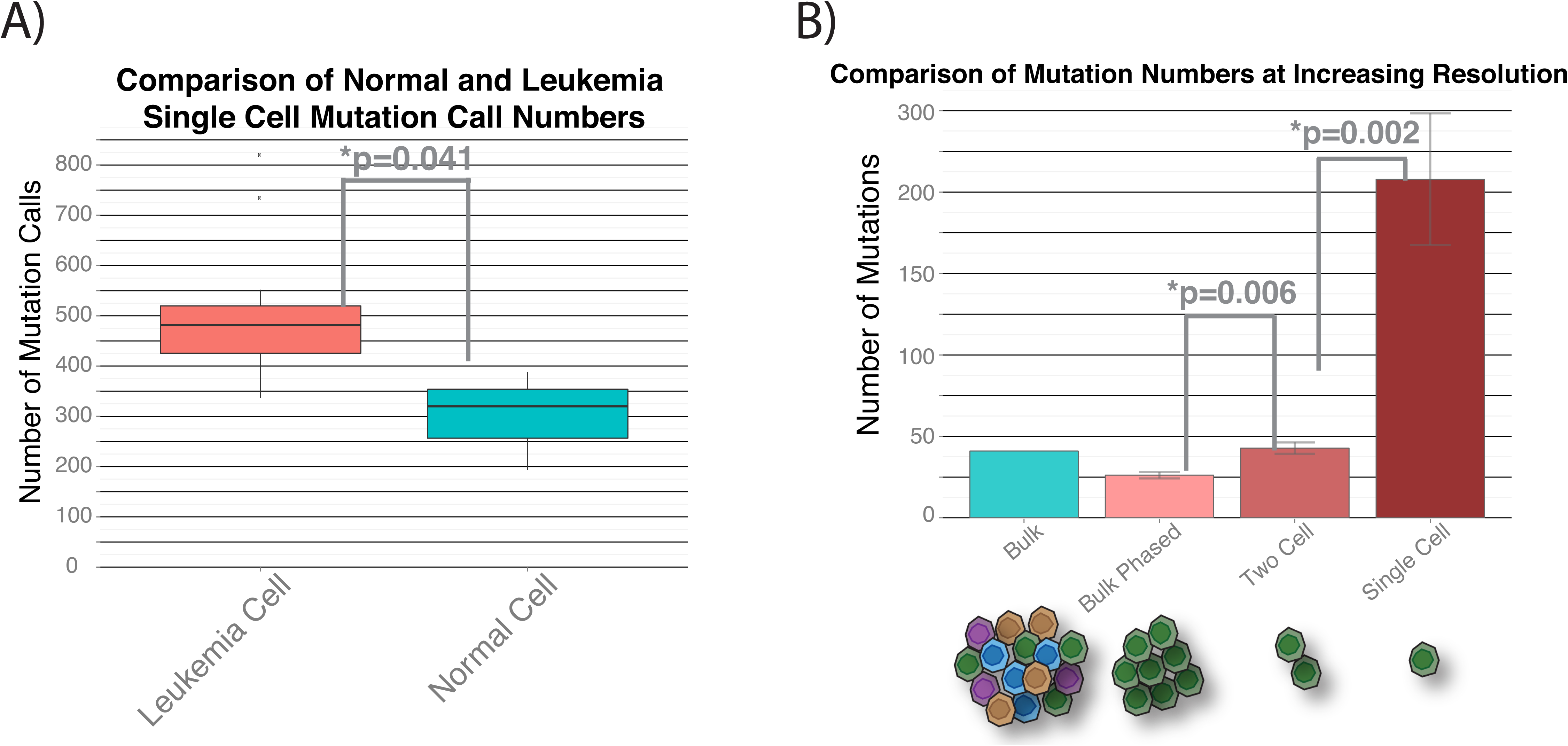
Estimating single-cell mutation rates. (A) Comparison of the number of exome mutation calls in leukemia cells to that in normal cells after removing germline SNVs. Subtracting the number of mutation calls in the normal cells provides an estimate of the mutations acquired by each leukemia cell. (B) Increasing the resolution of mutation calls identifies lower-frequency mutations. The increasing genetic diversity at higher-resolution measurements further supports the existence of much higher population genetic diversity.

We hypothesized that even more sensitive measurements would identify additional *RAS* mutations. We performed limited sample dilutions followed by error-corrected sequencing of patient SJETV075, hypothesizing that some *RAS* mutations that were just below our detection threshold would randomly be present at slightly higher frequencies in the dilute samples. By this approach, we identified 3 additional low-frequency *KRAS* mutations (G12R, A146T, L19F) for a total of 9 known activating *RAS* mutations in patient SJETV075. Our examination of all *RAS* mutations in this cohort revealed that of the 33 mutations identified, 31 were C-to-T or C-to-G changes. The 5 C-to-G changes were enriched for the APOBEC motif, but no similar enrichment was found in the C-to-T mutations, suggesting that they resulted from near-target APOBEC activity or replication-associated mutagenesis. This contrasts with lung cancer, in which approximately 60% of *KRAS* mutations are G12C or G12V, resulting from C-to-A substitutions^20^. Together, these observations further support the hypothesis that APOBEC activity and replication-associated mutagenesis are the underlying processes driving the evolution of *ETV6-RUNX1* ALL.

We then considered the variables that would result in such a large intra-patient activating *RAS* mutation burden. In our first model, we proposed that a large population of cells was at risk for rapid RAS-mediated expansion and underwent a widespread mutagenesis process, resulting in the concurrent outgrowth of multiple clones containing mutations conferring similar fitness. In our second model, we proposed that a smaller population was at risk for transformation, with a lower global mutation rate, but that *RAS* was a mutational hotspot, resulting in the acquisition of multiple *RAS* mutations during the period of leukemic transformation. If the *RAS* genes were mutational hotspots for *ETV6-RUNX1* ALL, we would expect to find additional benign mutations as passengers of the activating mutations. However, even our more sensitive approach to mutation detection, using both single-cell and error-corrected sequencing, found no additional somatic variants within *KRAS, NRAS*, or *HRAS*.

Taken together, these findings support the model in which a massive mutation burden develops within a relatively short time in a population at risk for transformation. To further support this model, we measured the mutation rate in single leukemia cells and used the normal cells from the same sample that had undergone whole-genome amplification in the same microfluidic chip to control for the background mutations and for amplification and sequencing errors. By this approach, we measured a mean of 208 coding mutations per cell that were above the background rate in the normal cells (Fig. 3A). Compared to our previous mutation estimation, this approach estimated greater genetic diversity, with each cell harboring a mean of 150 coding variants that were not detected by bulk sequencing or intraclonal exome sequencing (Fig. 3B).

### Simulation estimates the size and genetic diversity of leukemic populations

Given the large number of mutations per cell, we devised a model to estimate the population size in order to approximate the total genetic diversity across all the clonal populations at the time ALL is diagnosed. To accomplish this, we used those variables that could be estimated from the results of previous studies, along with our current measurements, to simulate *ETV6-RUNX1* ALL development. We know that the disease is initiated from a single cell by an *ETV6-RUNX1* translocation, as all cells harbor the same breakpoint^5^. Furthermore, each population acquires a mean of 12 deletions^9^. We also know that the disease is initiated *in utero* and develops over a mean of 4.7 years^21^; that lymphocyte precursors divide approximately every 11.9 days^22^, whereas primary leukemic cells divide much more frequently; and that replication errors occur in coding regions with a frequency of approximately once in every 300 human cell divisions^24^. From our results, we estimate that each cell had acquired approximately 200 coding SNVs, with many of them arising from APOBEC mutagenesis that occurred in a burst over a short time, randomly resulting in the 16 activating *RAS* mutations identified in our *ETV6-RUNX1* ALL cohort. Cells harboring *ETV6-RUNX1*, a recurrent deletion, and a *RAS* mutation have significantly increased rates of replication, resulting in the clinical diagnosis when children reach a total leukemic cell burden of approximately 1 ×10^11^ cells (Fig. 4A).

**Figure 4.**
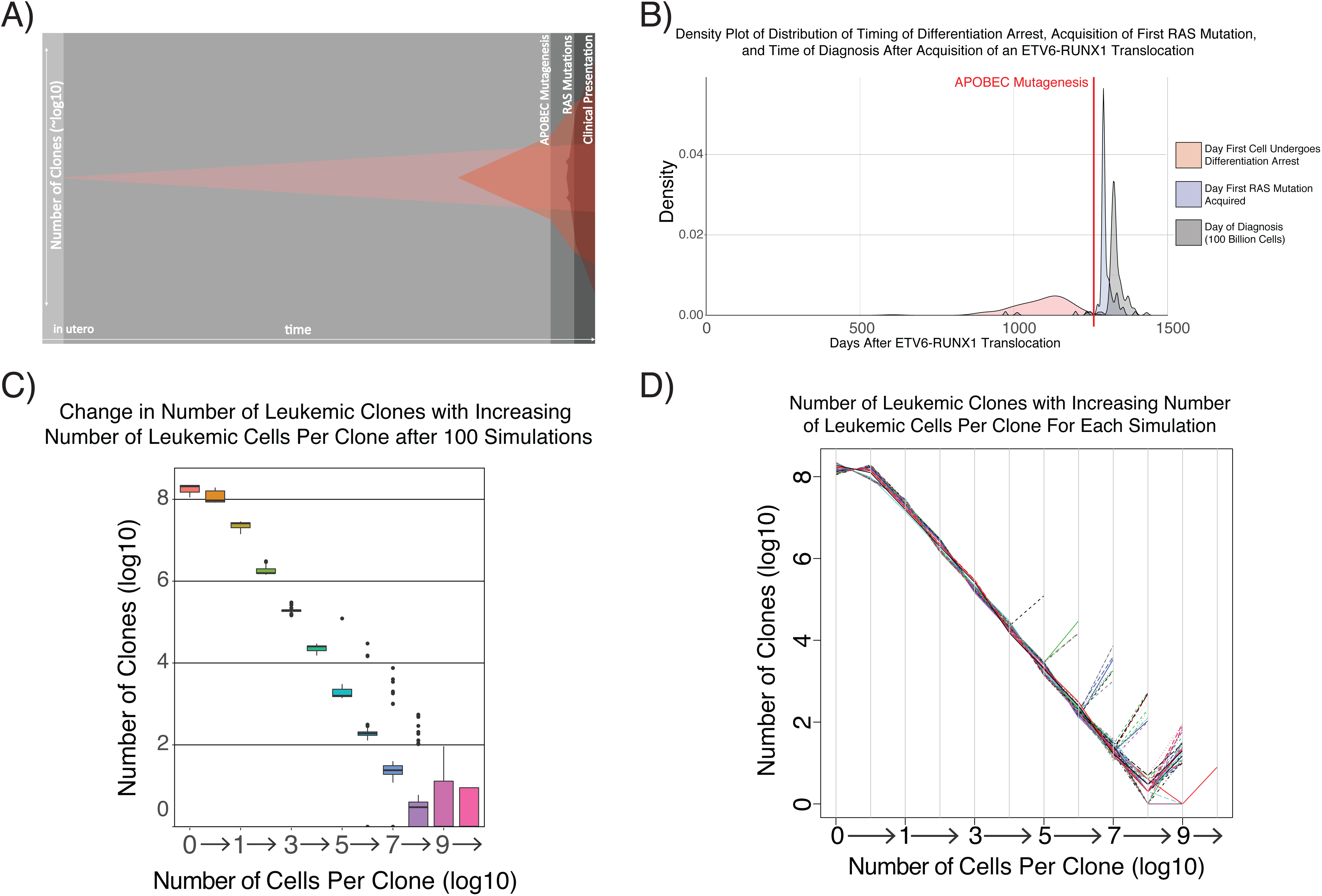
One hundred simulations of the development of *ETV6-RUNX1* ALL. (A) Overview of *ETV6-RUNX1* simulation in which a single cell with an *ETV6-RUNX1* translocation evolves over the years. Cells that acquire deletions that cause differentiation arrest are at risk for transformation by either replication or APOBEC mutations that create an activating *RAS* mutation. The cells expand until the subject acquires a total leukemic cell burden of approximately 1 ×10^11^. (B) The timing of differentiation arrest, APOBEC mutagenesis, appearance of the first activating *RAS* mutant clone, and clinical presentation of disease, using the described parameters, are similar to what is seen in patients. (C) When defining a clone based on the somatic coding mutation profile of each cell, there is an inverse correlation between the size and frequency of clones. The number of clones with the largest number of cells is more variable across simulations as a result of the timing and frequency of activating *RAS* mutations in cells that already harbor an *ETV6-RUNX1* translocation and a deletion that causes differentiation arrest. (D) Tracking the clone size and frequency for each of the 100 simulations shows the variance in the number of high-frequency clones across simulations.

Using those parameters, we initiated 100 simulations in which the mutation rates resulted in each cell acquiring a mean of 13 deletions and 229 SNVs by a mean of 28 days after a burst of APOBEC mutagenesis (**Supplementary Table 2**, Fig. 4B). Defining clones based on cells with the same somatic mutation profile in coding regions, we estimate that there were 330 million clones in total (Supplementary Table 2). However, most clones were created when the overall population size was high, making them rare (Fig. 4C). The number of high-frequency clones was variable and depended on the time and frequency of *RAS* mutation acquisition (Fig. 4D). We also found evidence that APOBEC mutagenesis and not replication errors caused most of the distinct *RAS* mutations and that the number of unique *RAS* mutations varied considerably between simulations (**Supplementary Fig. 3**). Taken together, these simulation results are consistent with our experimental data and support the assertion that there is massive population genetic diversity at the time a patient is diagnosed with ALL.

### Massive population diversity drives treatment dynamics in a patient sample

With such high population genetic diversity, we hypothesized that some clones already harbored mutations at diagnosis that altered their susceptibility to treatment. To test that hypothesis, we exposed leukemic cells to 5 standard ALL chemotherapy drugs (mercaptopurine, vincristine, prednisolone, daunorubicin, asparaginase) and used exome sequencing to evaluate the mutational composition after drug exposure. From this initial screen, we identified 537 putative mutations in at least one treatment. We then performed error-corrected sequencing of those sites in triplicate for each treatment to confirm that the mutations were pre-existing. We detected 224 specific base changes at the same location in more than one sample (**Supplemental Table 3**)—approximately 5 times as many mutations as the 41 that we identified in the initial bulk sequencing.

To identify resistant clones, we treated the cells with an increasing dose of each chemotherapy drug and looked for mutations exhibiting a dose-dependent increase in frequency. The control cells treated with no drug or DMSO, as well as the cells treated with the lowest dose of asparaginase, showed an increase in a distinct cluster of mutations. This cluster included the highest-frequency 146V *KRAS* mutation, along with 2 nonsynonymous *TP53* mutations (Fig. 5, cluster 7). Interestingly, 2 mutations (*FOLH1* R281H and *RGPD3* P816C) were strongly selected for in all control and treatment samples but were not detected in the diagnostic sample, suggesting they were selected for by the culture conditions. Treatment with low-dose mercaptopurine or higher doses of asparaginase resulted in reduced expansion of the *KRAS* A146V clone. This is consistent with the known pharmacokinetics of asparaginase, whereby increasing the dose after target saturation has no effect on cell killing^25^. Treatment with vincristine or dose level 2 of mercaptopurine further decreased the clone(s) in mutation cluster 7 and decreased the frequency of the highest-frequency mutations in cluster 1. Finally, exposure to prednisolone, the highest doses of mercaptopurine, or daunorubicin resulted in the greatest decrease in mutations in clusters 1 and 7, whereas mutations in clusters 2 through 4 increased in frequency as a result of these treatments. Treatment with the highest doses of daunorubicin killed all cellular populations. Taken together, these data show that the underlying genetic diversity when ALL is diagnosed does affect the treatment dynamics.

**Figure 5.**
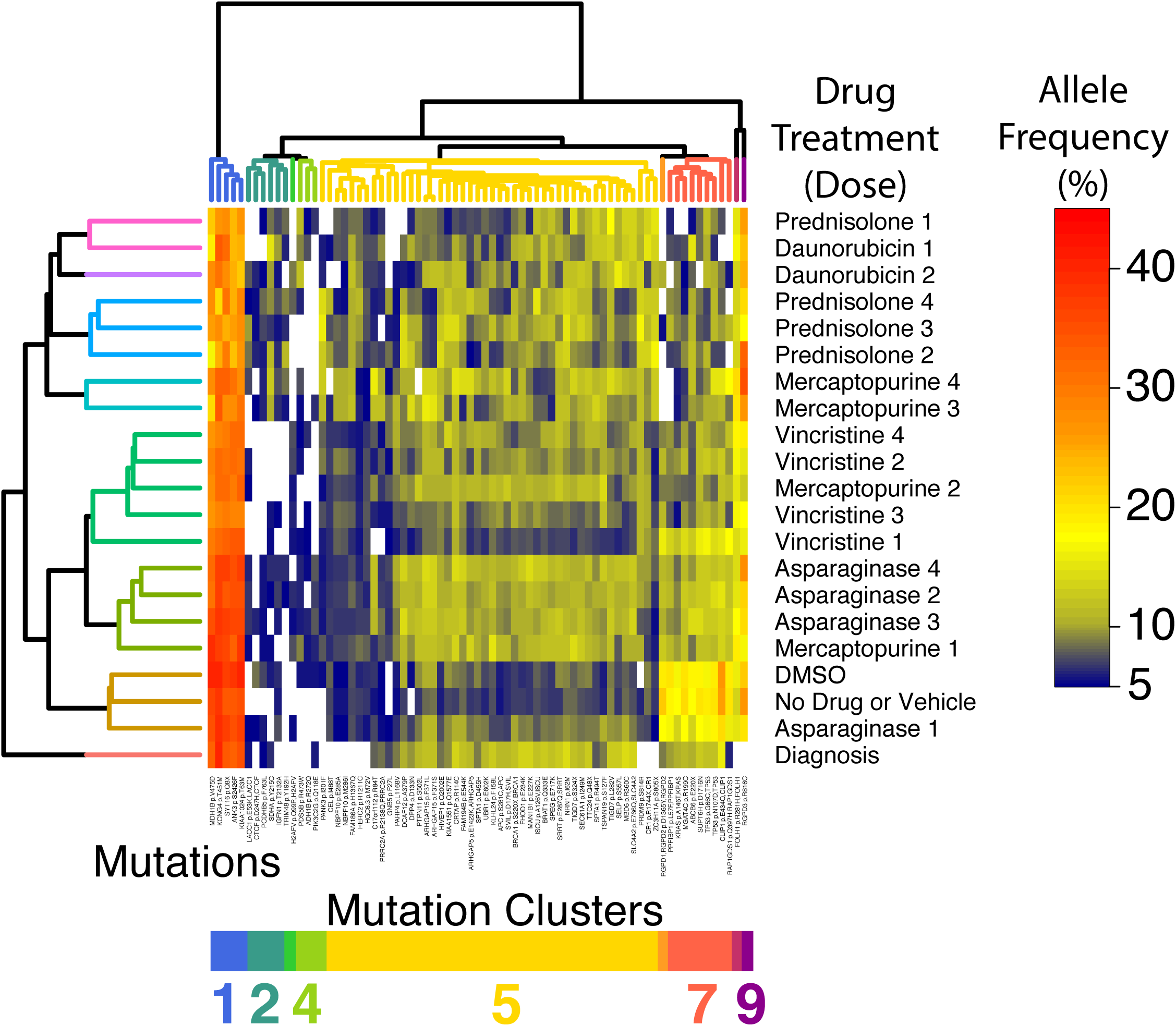
Differential sensitivity of leukemic populations to chemotherapy. Clusters of mutations showing patterns of response to drug treatment and dosage. Clones with cluster 7 mutations, which includes *KRAS* A146V and two *TP53* mutations, expanded without treatment or upon exposure to low-dose asparaginase when compared to the diagnostic sample that was not placed in culture. Mutations in clusters 8 and 9 were selected in all samples under the culture conditions used. Low-dose mercaptopurine or higher doses of asparaginase limited the expansion of mutation cluster 7. Exposure to vincristine or an increased dose of mercaptopurine further reduced mutation cluster 7. Higher doses of mercaptopurine, as well as exposure to prednisolone or daunorubicin, further decreased the frequency of mutations in clusters 1 and 7 while selecting for clones with mutations in clusters 2 through 4.

## Discussion

We have presented the results of a combination of error-corrected and single-cell sequencing of *ETV6-RUNX1* ALL cells collected at diagnosis in order to further resolve the temporal changes in clonal structures and mutational processes that occur during the development of the disease. The shift in the relative frequency of different types of cytosine mutation revealed an APOBEC mutagenesis pattern that decreases over time, suggesting that this process is important for disease progression but is not required for persistence or ongoing expansion of leukemic clones. We detected an unexpectedly high population mutation burden, which revealed differences in the treatment response among the clones that arose from the population that had previously undergone APOBEC and replication-mediated mutagenesis. This finding emphasizes the dynamic nature of leukemic evolution, as the relative importance of mutations required for cell survival shifts under distinct selection pressures as patients undergo treatment. We have integrated these findings with previous knowledge to create a new model of *ETV6-RUNX1* ALL, which is presented in Figure 6.

**Figure 6.**
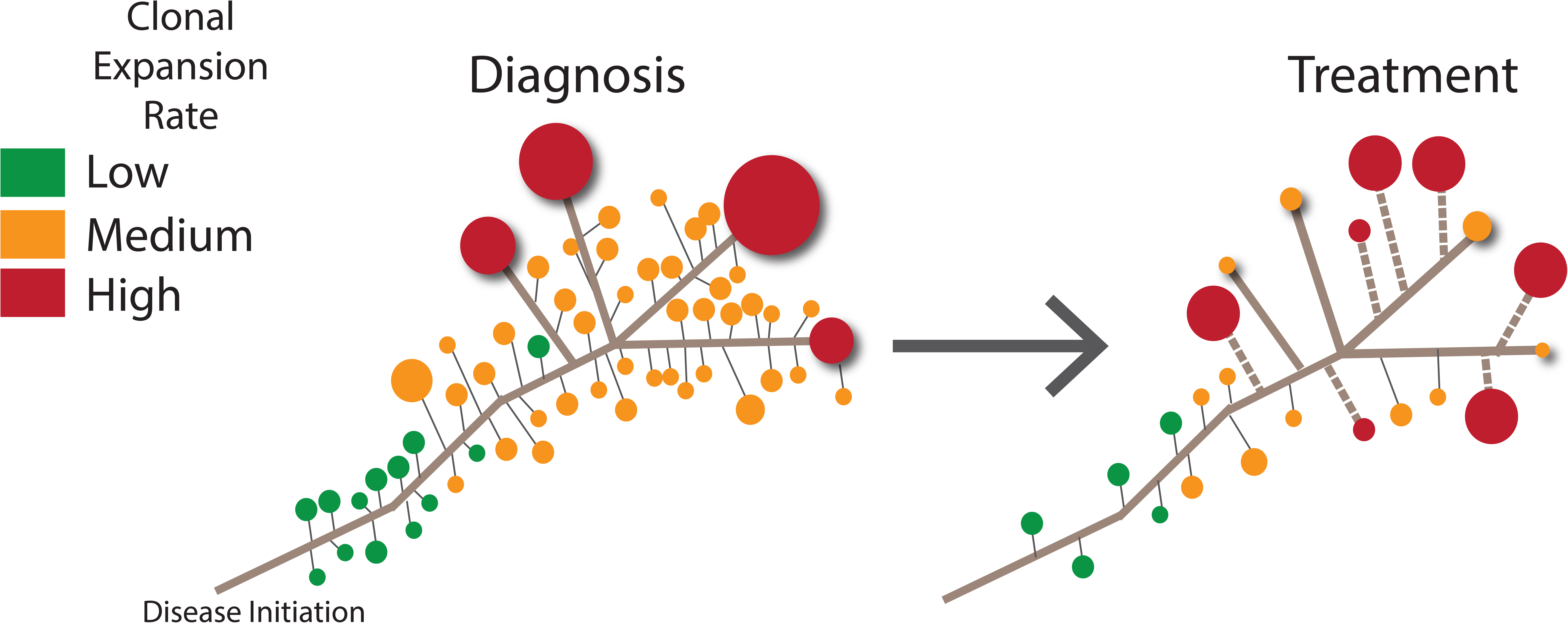
Model of *ETV6-RUNX1* evolution. A massive number of clonal populations are present at diagnosis, with a subset containing variants that allow them to become the most abundant (red). After selection pressures change during treatment, some of the high-frequency clones contract, whereas other clones with different somatic mutation profiles undergo positive selection (dashed line).

These new details of temporal shifts in the mutational processes of ALL in the period leading up to a patient's clinical presentation highlight how single-cell genomics can be used to trace back the mutational histories of tumors. The decrease in APOBEC-mediated mutagenesis during disease progression removes a major contributor to the global mutation rate in those patients. This change in the mutation rate may be one reason why ALL patients are cured at such high rates compared to patients with other tumor types in which the source of the mutagenesis, such as a mutation that decreases the fidelity of the DNA repair pathway, does not decrease over time^26^. From our results, it is unclear what effect ongoing exposure to mutagenic chemotherapy agents has on the induction of drug-resistant clones, but the effect could be significant in pediatric patients with ALL, most of whom undergo treatment for 2 to 3 years^27^.

Mutations that are considered driving lesions, such as variants in *KRAS* or *TP53*, were not dominant at diagnosis or after undergoing selection during drug treatment, suggesting that combinations of mutations mediate the clinical behaviors of clones. We also observed the rise of clusters of mutations, but it is unclear how many distinct clonal populations those clusters represent. This is an area where higher-resolution studies using single-cell sequencing to determine the co-occurrence patterns of mutations will provide additional insights into drug resistance. The putative high number and differential drug sensitivity of leukemic clones also provide insights into the need for combination therapy to cure children with ALL. Although the sheer genetic diversity of the population presents new challenges for the design of targeted treatment strategies, the presence of massive population genetic diversity in ALL also provides an opportunity to probe those samples to learn why we can already overcome the pre-existent resistance in most patients with ALL. Focusing on specific mutations that are selected for by particular drugs could yield mechanistic insights into drug-specific resistance and provide a new rationale for choosing drug combinations.

In summary, we have provided new insights into the development *of ETV6-RUNX1* ALL and its resistance to treatment by studying population genetics. Our findings underscore the importance of studying the mutation burden, size, and mutation rate of the population as a whole, not just the highest-frequency variants detected by standard bulk sequencing, when trying to understand and predict treatment response. Greater genetic diversity has also recently been reported in other premalignant and malignant states^28,29^, suggesting that the study of cancer population genetics is important for understanding most tumor types.

Together, these studies have revealed new layers of complexity in leukemic evolution that need to be fully understood in order to more effectively eradicate premalignant and malignant ALL cell populations with less treatment-related toxicity, and they have provided a framework for studying intra-tumor evolution in a wide range of malignant neoplasms.

## Online Methods

### Single-cell exome sequencing and mutation calling

Amplified DNA from patient 4 that had undergone single-cell isolation and whole-genome amplification using the Fluidigm C1 System as previously described^10^ was used for library construction and exome capture with the Nextera Rapid Capture Exome Kit (Illumina), used in accordance with the manufacturer's instructions. Exome-enriched libraries then underwent sequencing using 2 xx100 reads on 4 flow cells of a HiSeq 2000 or 2500 Sequencing System (Illumina). Adapters were trimmed from each of the cells by using Trimmomatic (ILLUMINACLIP:nextera_adapters.fa:2:30:10 TRAILING:25 LEADING:25 SLIDINGWINDOW:4:20 MINLEN:30), followed by alignment with BWA using default parameters. Duplicates were marked using Picard (https://broadinstitute.github.io/picard), and local realignment followed by base score recalibration was performed using GATK (https://software.broadinstitute.org/gatk). We then called variants by using GATK and followed this with filtering using the parameter “QD < 2.0 || FS > 60.0 | | MQ < 40.0 | | HaplotypeScore >13.0 | | MQRankSum< −12.5 | | ReadPosRankSum< −8.0”. On-target coverage was calculated with Picard HsMetrics; this was repeated after subsampling for an increasing number of reads using custom bash scripts. Custom bash scripts were also used to identify locations that had the same mutation called in more than one cell. Germline SNP locations identified by bulk sequencing were then filtered out, after which locations that were identified in any of the normal single cells were removed.

### Error-corrected sequencing

Adapters with unique identifiers were prepared as previously described. Aliquots of 250 or 500 ng of genomic DNA then underwent 30 min of chemical fragmentation and standard library preparation by using the KAPA HyperPlus Kit (Kapa Biosystems) with adapters that contained unique molecular identifiers as described^30^, using 3 μg of adapter per reaction (a 10:1 molar ratio). PCR amplification and hybrid capture were performed as previously described^31^. Sequencing was performed using MiSeq V2 chemistry, using 2 × 150-bp PE reads. We then trimmed the sequences to 125 bp with Trimmomatic and placed the unique molecular identifiers into the header by using the script tag_to _header.py^30^. Reads were aligned using BWA ALN with standard parameters. We sorted and indexed using Picard then performed consensus calling by using ConsensusMaker.py with parameters–minmem 3,–cutoff 0.8, and --Ncutoff 0.7. Unmapped reads were removed with SAMtools (http://samtools.sourceforge.net/), then local realignment was performed using GATK before creating an mpileup file. Normal and Tumor mpileup files were then compared using VarScan Somatic, with somatic mutations requiring a *P*-value of less than 10^−4^, as computed using Fisher's exact test by VarScan (http://varscan.sourceforge.net). We also required the germline sample to have fewer than 5 reads and that no more than 90% of variant reads were on the same DNA strand. Variants underwent RefSeq annotation with ANNOVAR.

### Estimation of mutation rates

To estimate the mutation rate, we downsampled each of the files to 70 million reads. We then created marked, realigned, and base score–recalibrated BAM files as described above. This was followed by further variant calling and filtering using the GATK filtering parameters detailed above. We then subtracted those sites that were found in the bulk germline sequencing. To subtract the background error rate due to amplification errors, the somatic mosaicism rates in normal cells, and mutation miscalls, we subtracted the mean mutation rate in the 3 normal cells from that of each of the single tumor cells and plotted the distribution of the mutations rates.

### Simulation

We designed a computational model to investigate when RAG-mediated deletions and *RAS* mutations occur, the clonal burden, and the timing of disease onset. The model is initiated with a single cell with an *ETV6-RUNX1* translocation that is capable of differentiation and divides every 12 days, defined as cell type 0. At each time point, cells may gain a deletion or, if dividing or if APOBEC mutagenesis is active, an SNV. Type 1 cells are created when type 0 cells gain a specific RAG-mediated deletion, which occurs with probability *pRAG_DA.* Type 1 cells are considered to have differentiation arrest, but they divide at the same rate as type 0 cells. Type 2 cells are created when type 1 cells have a mutation within *RAS*, which occurs with probability *pMut_IP.* Type 2 cells have differentiation arrest as well as increased proliferation, duplicating every day. The number of cells within each cell type with a specific amount of mutations and deletions are tracked. One time point is equivalent to 1 day, and each cell is stochastically sampled.

For each cell of a given type, a random number is generated from 1 to 1/*pDel*, where *pDel* is the probability of gaining a deletion. A deletion occurs within cells for which this random number equals 1. A second random number is generated from 1: 1/*pRAG_DA* for each type 0 cell that acquires a deletion. Any cell in which this second number is 1 contains a deletion that causes it to transform into a type 1 cell. This same process is carried out when calculating the mutational burden as well as the transformation of type 2 cells into type 3 cells.

APOBEC mutagenesis starts on a fixed day, if cell type 3 has not been generated, and continues for 2 days after cell type 3 has formed. Mutations generated from APOBEC are calculated in the same way as previously described, with the exception that every cell within cell types 1 and 2 undergoes APOBEC mutagenesis and the number of mutations caused by APOBEC within a given cell is determined by randomly sampling from *1:Burst*. The simulation completes when cell type 3 has reached 1 ×10^11^ cells. We estimate the number of cells per clone by the timing of the newly acquired mutation and replication rate, accounting for the branching of cells such that the sum of all cells equals the number of cells of a given type.

Simulations were run using our new R package, which is called RepALL and is available at https://github.com/mjdm/RepALL. We ran the simulation with the following default parameters:

Probability of gaining a RAG-mediated deletion, *pRAG:* .008
Probability of causing differentiation arrest, given a deletion (Type 0), *pRAG_DA:* 2e-7
Probability of gaining a mutation due to replication error, *pBGMut:* 0.003̅
APOBEC mutagenesis start day: 1290
APOBEC duration after leukemic cell-type initiation: 3 days
Maximum number of APOBEC mutations generated per cell per day, *Burst:* 75
Probability of gaining a *RAS* mutation, given a mutation (Type 1), *pMut*_IP: 16/(3e9*.02) = 2.6̅e-7

### Primary cell culture and drug treatment

Primary samples were from patients that had provided consent in studies approved by the St. Jude IRB. At the time of sample collection, mononuclear cells were isolated using Ficoll-Paque (GE Life Sciences) followed by cryopreservation. One vial of cells from each patient was thawed slowly using the ThawSTAR system (MedCision), and the cells were placed in culture under the conditions previously described^32^. For the limited dilution experiment, 750,000 cells were plated in each well of a 12-well plate and were grown in culture for 3 weeks. For drug treatments, the drugs and 350,000 cells were plated in each well of a 24-well plate. In both experiments, the medium was changed twice weekly by carefully removing half of it and replacing it with fresh medium. The replacement medium included a 2× drug concentration if the cell sample was undergoing chemotherapy exposure. All drugs were purchased from Sigma-Aldrich, and the concentration ranges were based on solubility limits and previously published data^32^. The drugs used were mercaptopurine (500, 250, 125, and 62.5 μg/mL ConsensusMaker.py and 90 μg/mL), vincristine (810, 162, 32.4, and 6.5 μg/mL), daunorubicin (31, 6.2, 1.2, and 0.2 μg/mL), and asparaginase (19, 9.5, 4.8, and 2.4 μg/mL). Live cells were isolated by using a dead cell removal kit (Miltenyl). DNA was extracted using a DNA Universal Kit (Zymo Research), and libraries were prepared using the HyperPlus Kit (Kapa Biosciences). Exome or custom capture was performed using oligonucleotides and the standard protocol from Integrated DNA Technologies. Quality trimming, alignment, and mutation calling were performed using the pipeline outlined above.

## Acknowledgements

The authors would like to acknowledge Stephen Quake, Jinghui Zhang, and Jim Downing for their constructive advice on the project. In addition, the authors would like to thank the members of the Pediatric Cancer Genome Project, especially Charles Mullighan and Ching-Hon Pui who lead the Hematological Malignancies Program. V.G., Y.I., J.E., and C.G. are supported by ALSAC. C.G. is also supported by the Burroughs Wellcome Fund, Leukemia and Lymphoma Society, Hyundai Hope on Wheels, and the American Society of Hematology.

The data are available in the short read archive accession ID.

## Author contributions

C.G., V.G., and M.M. designed research; V.G., J.E., M.M., and C.G. performed research; C.G., M.M., and I.D. contributed new reagents/analytic tools; C.G., and M.M. analyzed data; and C.G., V.G., M.M., and I.D. wrote the paper.

## SUPPLEMENTARY MATERIALS

**Supplementary Table 1.** List of ALL hotspot mutation locations in the error-corrected sequencing capture panel.

**Supplementary Table 2.** List of recurrent mutations detected in patient SJETV077.

**Supplementary Table 3.** Statistics from the simulation of the development of *ETV6-RUNX1* ALL.

**Supplementary Figure 1.** Saturation of sequencing coverage at increasing depth. (A) Exome sequencing of bulk samples reached a saturating coverage breadth of 94% at 40 million reads. (B) Single cells reached a saturating coverage breadth of 82% at 60 million reads.

**Supplementary Figure 2.** Distribution of duplicate numbers in error-corrected sequencing. Unique molecular identifier family size distributions for (A) germline and (B) leukemia samples.

**Supplementary Figure 3.** Estimating the relative contribution of APOBEC and replication mutagenesis to activating *RAS* mutations. There is significant variability in the number of activating *RAS* mutations produced by each simulation. Most of the activating *RAS* mutations in the simulations were produced by APOBEC (A) rather than by replication-mediated (B) mutations.

